# Cortical and raphe GABA_A_, AMPA receptors and glial GLT-1 glutamate transporter contribute to the sustained antidepressant activity of ketamine

**DOI:** 10.1101/2020.03.04.976308

**Authors:** Thu Ha Pham, Céline Defaix, Thi Mai Loan Nguyen, Indira Mendez-David, Laurent Tritschler, Denis J David, Alain M. Gardier

## Abstract

At sub-anaesthetic doses, ketamine, a non competitive N-methyl-d-aspartate (NMDA) receptor antagonist, has demonstrated remarkable and rapid antidepressant (AD) efficacy in patients with treatment-resistant depression (TRD). However, its mechanism of action of ketamine is not fully understood. Since comorbid depression and anxiety disorders often occur, GABAergic/inhibitory and glutamatergic/excitatory drug treatments may be co-administered in these patients. Information regarding this combination is critical to establish efficacy or treatment restrictions to maximize translation from animal models to TRD patients, effectiveness and safety. To assess the specific role of excitatory/inhibitory neurotransmission in the medial prefrontal cortex-raphe nuclei (mPFC-DRN) circuit in the sustained antidepressant-like activity (AD) of ketamine (at t24h post dose), AMPA-R antagonist (intra-DRN) and GABA_A_-R agonist (intra-mPFC) were co-administered with ketamine (intra-mPFC). Twenty-four hours later, responses in the forced swim test (FST) and neurochemical consequences on extracellular mPFC glutamate, GABA and 5-HT levels were measured in BALB/cJ mice. Intra-DRN NBQX prevented the sustained AD-like activity of ketamine evidenced by decreases in FST swimming duration and blunted cortical 5-HT_ext_ and Glu_ext._ Intra-mPFC muscimol blocked ketamine AD-like activity and its effects on cortical 5-HT_ext_. Moreover, a selective glutamate transporter GLT-1 inhibitor, dihydrokainic acid (DHK) locally perfused into the mPFC produced an AD-like activity at t24h associated with robust increases in mPFC 5-HT_ext_, Glu_ext_ and GABA_ext_. Thus, the sustained AD-like activity of ketamine is triggered by AMPA-R activation in the DRN and 5-HT - glutamate release in the mPFC, but limited by GABA_A_-R activation - GABA release in the mPFC. The local blockade of GLT-1 in the mPFC also mimics the rapid responses of ketamine, thus highlighting the role of neuronal-glial adaptation in these effects. These results also suggests the need to test for the concomitant prescription of ketamine and BZD to see whether its sustained antidepressant activity is maintained in TRD patients.

## 1. Introduction

Ketamine, a non-competitive antagonist of the N-methyl-D-aspartate receptor of glutamate (NMDA-R) displays an antidepressant efficacy in treatment-resistant depression (TRD) (Berman et al., 2000; Zarate et al., 2006). Since this discovery, multiple randomized clinical trials confirmed this efficacy (see meta-analysis reviews: Caddy et al., 2014; Fond et al., 2014; Newport et al., 2015; Xu et al., 2016). The search for a new treatment of TRD is important because ≈30% of depressed patients are affected by refractory depression (Rush et al., 2006). Furthermore, the delayed onset of action of classical antidepressant drugs (i.e., 4 to 6 weeks for selective serotonin reuptake inhibitor, SSRI) makes the rapid ketamine action (less than 24h in human and animals) of great value, but we need more details about its mechanism of action.

Comorbid depression and anxiety disorders occur in up to 25% of patients (Tiller, 2013). For example, chronic neuropathic pain often leads to anxiety and depression disorders (Sellmeijer et al., 2018). Both disorders require appropriate treatment, e.g., an antidepressant drug for depression, and a benzodiazepine (BZD) for anxiety. In addition, a number of GABAergic (BZD) and anti-glutamatergic treatments are used as adjunctive therapy in TRD (Frye et al., 2015). Thus, it is especially important to know whether or not ketamine and a BZD (an agonist of GABA_A_ receptor) can be associated. It was recently shown that concomitant BZD use attenuated ketamine response (Frye et al., 2015). Such information is critical to maximize translation from animal models to TRD patients, effectiveness and safety.

Animal models of anxiety/depression contribute robustly to study ketamine’s mechanism of action (Pham et al., 2019). 80% of medial prefrontal cortex (mPFC) neurons are excitatory and the mPFC contains a dense network of glutamate releasing nerve terminals (Gasull-Camos et al., 2017) that project to the dorsal raphe nucleus (DRN) (Amat et al., 2016). In addition, NMDA-R, the main target of ketamine, is widely expressed in this brain region (Murray et al., 2000; Kamiyama et al., 2011; Sanz-Clemente et al., 2013). Artigas’s group demonstrated that 5-HT release in the mPFCx depends on the excitatory glutamatergic transmission (Lopez-Gil et al., 2012). Ketamine triggers a cascade of neuronal adaptation involving the mammalian target of rapamycin (mTOR) pathway in the mPFC and activates synaptogenesis in an α-amino-3-hyroxy-5-methyl-4-isoxazolepropionic-acid-receptor- (AMPA-R)-dependent manner (Li et al., 2010; Duman et al., 2016, 2019). Upregulation of AMPA-R synaptic expression has been described in rodents within 24 hours after ketamine treatment (Zanos et al., 2016). Furthermore, recent evidence suggests that ketamine requires an activation of AMPA-R to exert its antidepressant-like activity since NBQX, an AMPA-R antagonist, blocked ketamine responses in behavioral tests in rodents (Koike & Chaki, 2014; Koike et al., 2011; Li et al., 2010; Li et al., 2011; Maeng et al., 2008; Pham et al., 2018b). However, the targets by which ketamine produces glutamate bursts that trigger the fast and sustained (at t24h post dose) antidepressant-like activity of ketamine remain unclear (Fuchikami et al., 2015).

We recently described a positive correlation between mPFC 5-HT neurotransmission and ketamine-induced antidepressant-like activity in the forced swim test (FST) (Pham et al., 2018a, 2018b). The increased swimming duration in the FST is consistent with an antidepressant-mediated increase in serotonergic neurotransmission (Cryan et al., 2002). However, the origin of this increase in 5-HT release still remains questionable. The cell bodies of 5-HT neurons are located in the DRN, a region receiving dense glutamatergic projections from the mPFC (Fukumoto et al., 2016). A systemic administration of ketamine increases c-Fos immunoreactivity in DRN 5-HT neurons, which were blocked by NBQX microinjection into the mPFC (Fukumoto et al., 2016) suggesting that the activation of these neurons modulated by the mPFC could contribute to ketamine mechanism of action. Still, information about the influence of the mPFC on the firing activity of DRN 5-HT neurones and vice versa in ketamine-induced antidepressant-like activities are missing.

Here we assessed the behavioral and neurochemical effects of ketamine by coupling microdialysis in the mPFC and FST in BALBc/J mice. In a pharmacological approach, we used a pre-treatment with AMPA-R antagonist (intra-DRN NBQX: Nishitani et al., 2014), GABA_A_-R agonist (intra-mPFC muscimol: Amat et al., 2016) administered thirty minutes before ketamine to study the specific role of excitatory and inhibitory neurotransmission in the mPFC-DRN circuit. Extracellular levels of glutamate, GABA and 5-HT (Glu_ext_, GABA_ext_, and 5-HT_ext_, respectively) were examined in the mPFC. We also investigated neurochemical and behavioral consequences of glutamate transporter GLUT-1 (or EAAT2) blockade after intra-mPFC perfusion of dihydrokainic acic (DHK), a selective inhibitor of GLT-1, present in astrocytes (Gasull-Camos et al., 2017).

## 2. Materials and methods

### 2.1. Animals

Male BALB/cJ mice (9-12-weeks old) weighing 23-25g at the beginning of the experiments were purchased from Janvier Labs (Le Genest-Saint-Isle). The BALB/cJ strain of mice was chosen for its highly anxious phenotype (Dulawa et al., 2004; Holick et al., 2008; Calcagno and Invernizzi, 2010.). They were housed in groups of four in a temperature (21 ± 1°C) controlled room with a 12 h light: 12 h dark cycle (lights on at 06:00 h). Food and water were available ad libitum except during behavioral observations. Particular efforts were made to minimize the number of mice used in the experiments. Protocols were approved by the Institutional Animal Care and Use Committee in France (Council directive # 87-848, October 19, 1987, “Ministère de l’Agriculture et de la Forêt, Service Vétérinaire de la Santé et de la Protection Animale, permissions # 92-196” to A.M.G.) as well as with the European directive 2010/63/EU.

### 2.2. Drugs and treatments

Ketamine (2 nmol) purchased from Sigma-Aldrich (Saint-Quentin Fallavier, France) was dissolved in artificial cerebrospinal fluid (aCSF). Microdialysis samples and the FST were performed 24 h later, a time to measure ketamine sustained antidepressant-like activity. Both the FST and microdialysis technique have been performed in the same mice. In addition, this time point was chosen in order to avoid ketamine-induced hyperlocomotion and the psychotomimetic effects, which are observed in rodents when behavioral tests are performed immediately after an acute injection (Koike et al., 2013; Li et al., 2010). Drug doses and pre-treatment times were based on previous studies (Iijima et al., 2012; Koike et al., 2013; Li et al., 2010; Liu et al., 2012; Zanos et al., 2015; Pham et al., 2018b).

First, the AMPA-R antagonist, NBQX (0.25 nmol) was perfused into the DRN (NBQX disodium salt purchased from Tocris Bioscience, Lille, France). This dose was chosen based on previous studies in rodents (Lopez-Gil et al., 2007; Fukumoto et al., 2016). Second, the GABA_A_-R agonist, muscimol (8 nmol) was injected into the mPFC (Sigma-Aldrich, Saint-Quentin Fallavier, France), according to Amat et al., (2016). Third, the inhibitor of GLUT-1 glutamatergic transporter, dihyrokainic acid (DHK, Tocris Bioscience, Lille, France) was dissolved in the aCSF and perfused at 5 mM for 2.5 hours according to Gasull-Camos et al., (2017). The swimming duration in the FST was measured in these mice when mPFC dialysates were collected as shown in protocols (Figure 1A – 3A). Bilateral injections of ketamine, muscimol and DHK were used in the present study.

**Fig. 1.**
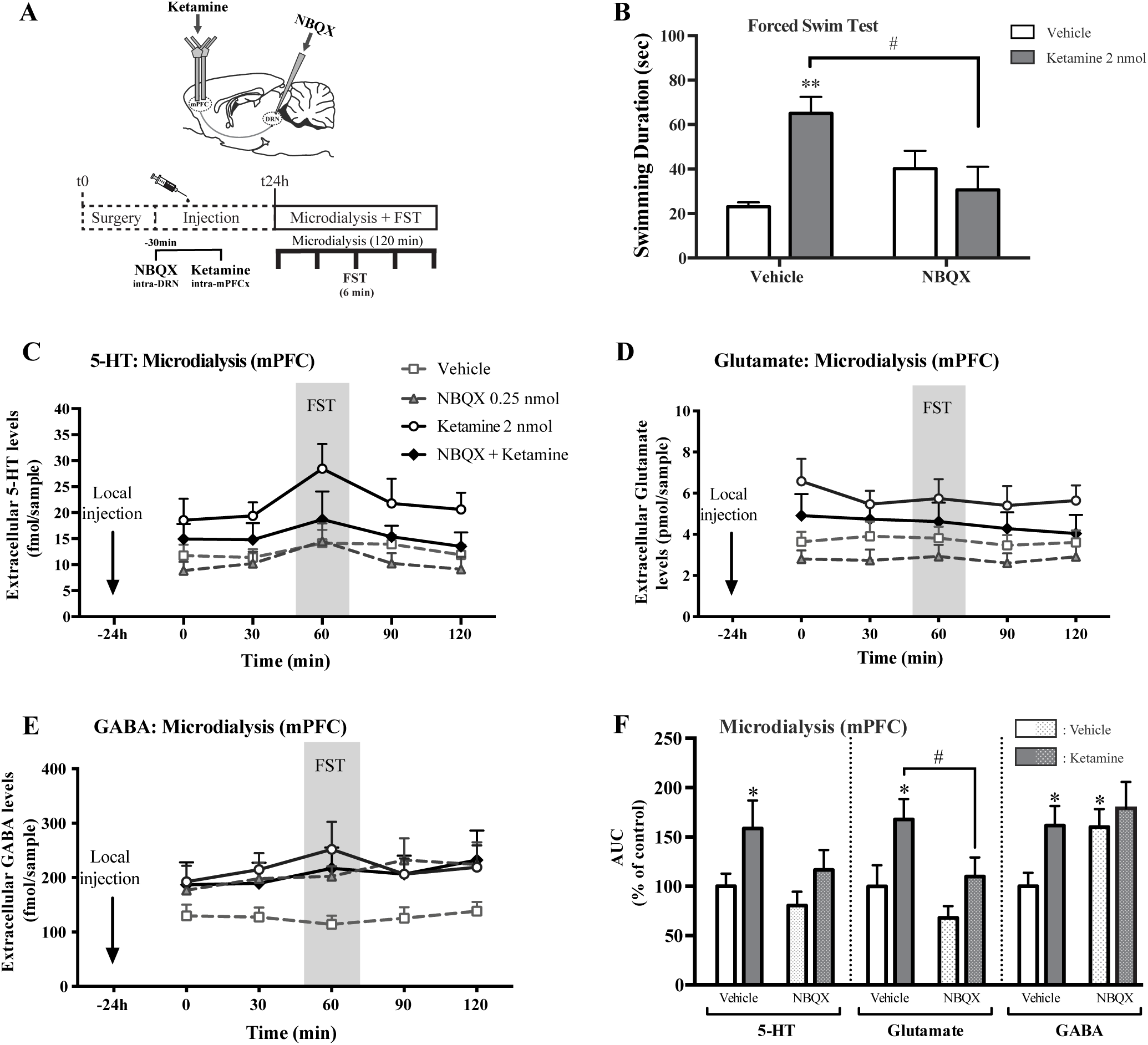
AMPA-R antagonist NBQX-pre-treatment into the DRN blocks the sustained antidepressant-like activity of ketamine and its releasing effects on 5-HT-glutamate-GABA in the mPFC. (A) Experimental protocol: after the surgery, NBQX 0.25 nmol was injected locally intra-DRN at 30 min prior to ketamine 2 nmol intra-mPFC. On the next day (t24h), the tests were performed in the same mice. The gray area in Fig. 1C, 1D and 1E indicates the duration of the FST (i.e., 6 min) performed during the collection of microdialysis samples. All dialysates were analyzed for extracellular 5-HT, Glu and GABA levels. (B) NBQX blocked ketamine effects on the swimming duration in the FST and (C) blunted ketamine-induced increase in 5-HT levels in the mPFC as shown in the time course t0-120 min. (D) and (E) effects of NBQX on the time course of ketamine-induced increase in Glu and GABA extracellular levels in the mPFC. (F) The AUC values were calculated for the amount of 5-HT, Glu and GABA outflow collected during 0-120 min, and expressed as percentage of control group. *p<0.05 vs Vehicle-treated group. #p<0.05 vs ketamine-treated group group (two-way ANOVA). Data are presented as means ± S.E.M (n=5 mice per group).

**Fig. 2.**
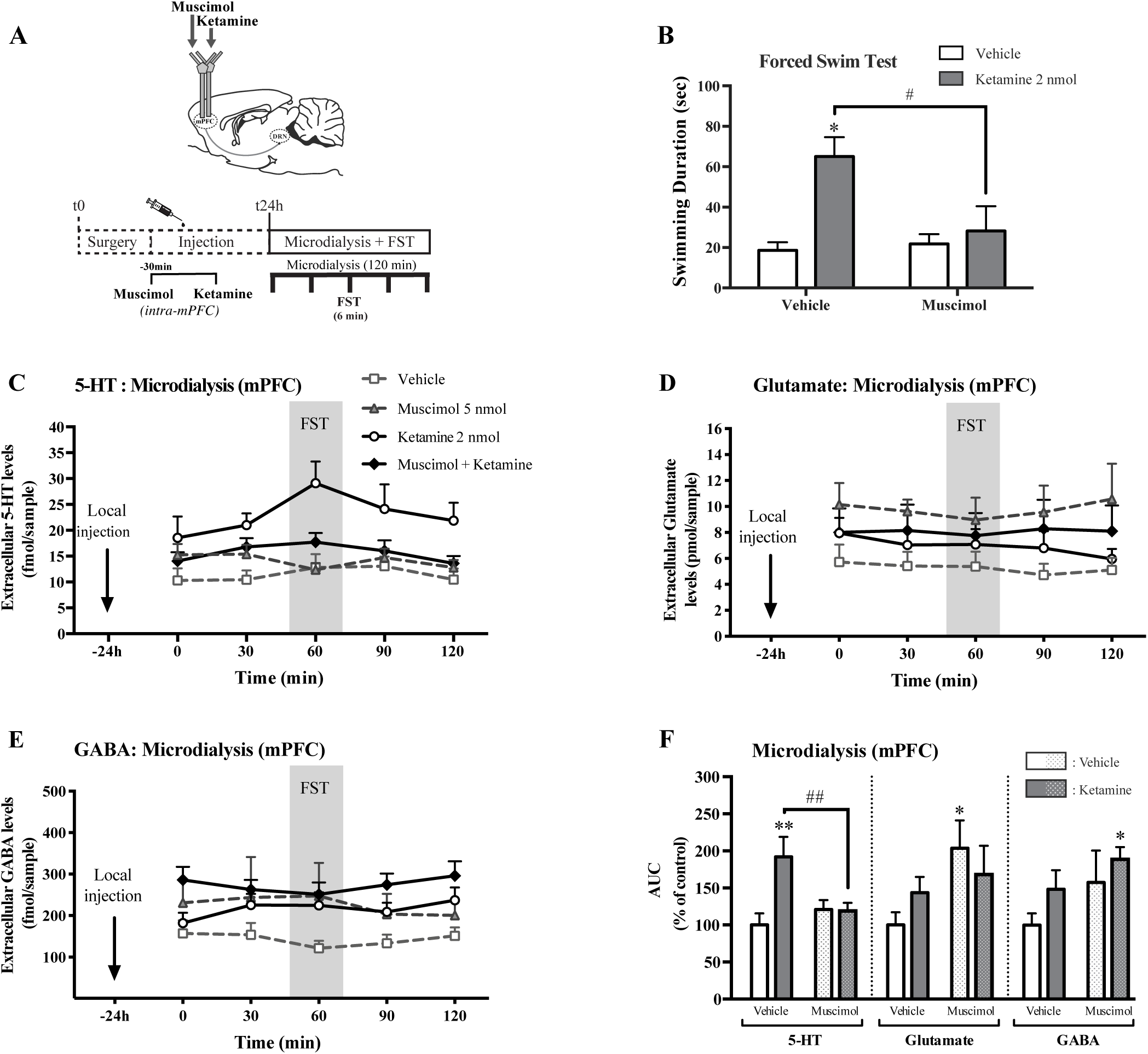
GABA_A_-R agonist muscimol pre-treatment in the mPFC decreases antidepressant-like activity and 5-HT releasing property of ketamine. (A) Experimental protocol: after the surgery, muscimol (2.5 nmol per side) was injected bilaterally intra-mPFC at 30 min prior to ketamine (1 nmol per side) injected in the same site. On the next day (t24h), the tests were performed in the same mice. The gray area in Fig. 2C, 2D and 2E indicates the duration of the FST (i.e., 6 min) performed during the collection of microdialysis samples. All dialysated were analyzed for extracellular 5-HT, Glu and GABA levels. (B) Muscimol blocked ketamine effects on the swimming duration in the FST and (C) that on 5-HT levels in the mPFC as shown in the time course t0-120 min. (D) Muscimol itself increased the extracellular Glu levels in the mPFC *versus* vehicle-treated group. (E) Muscimol did not modify ketamine-induced effects on extracellular GABA levels. (F) The AUC values were calculated for the amount of 5-HT, Glu and GABA outflow collected during 0-120 min, and expressed as percentage of control group. *p<0.05; **p<0.01 vs Vehicle-treated group. ##p<0.01 vs ketamine-treated group (two-way ANOVA). Data are presented as means ± S.E.M (n=4-5 mice per group).

**Fig. 3.**
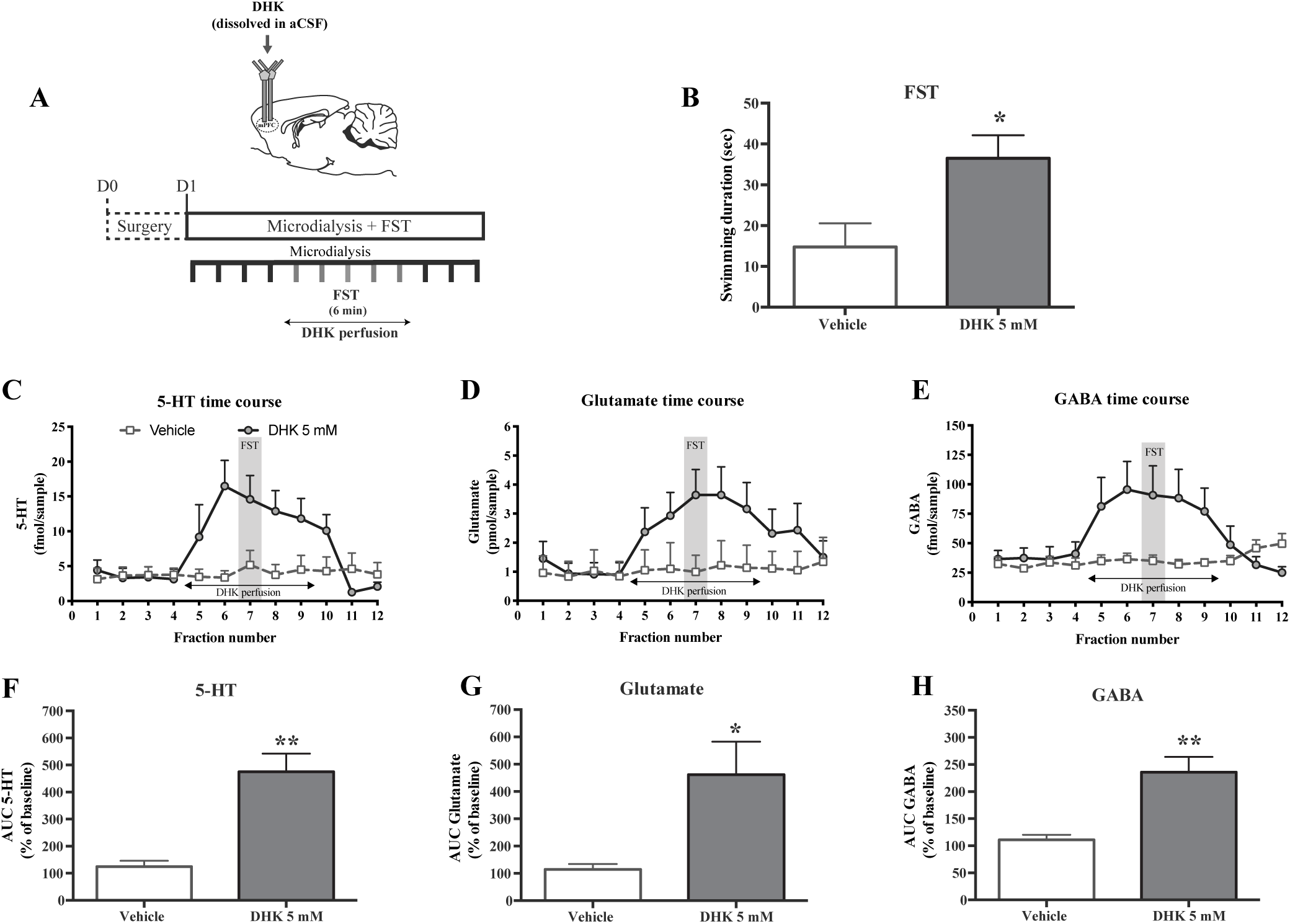
*Intra-mPFC DHK, a glial glutamate transporter GLT-1 inhibitor, mimics the sustained antidepressant-like effect and neurochemical effects of ketamine* t. (A) Experimental protocol: after achieving 4 points of baseline, DHK 5 mM (dissolved in aCSF) was perfused during 2.5 hours (equal 5 collecting points of dialysates). (B) DHK induced an increase of swimming duration in the FST. (C, D, E) Time course of extracellular 5-HT, glutamate and GABA levels in the mPFC, respectively, following local DHK injection. The gray area indicates the duration of the FST (i.e., 6 min) during the collection of microdialysis samples. Data are presented as fmol/sample for extracellular 5-HT and GABA levels, and as pmol/sample for glutamate levels. (F, G, H) DHK induced an increase in the three neurotransmitters. The AUC values were calculated for the amount of 5-HT, glutamate, and GABA, respectively, and expressed as percentage of baseline levels. * p<0.05, ** p<0.01 vs Vehicle-treated group. Data are presented as means ± S.E.M (n=4-6 mice per group).

### 2.3. Forced Swim Test (FST)

The mouse forced swim test procedure (FST) is used for antidepressant drugs screening. Swimming, climbing and immobility durations were previously distinguished from each other in BALB/cJ mice (Dulawa et al., 2004; Holick et al., 2008). Swimming behavior relies on the serotonergic system, and climbing behavior on the noradrenergic system in the mouse (Holick et al., 2008). Mice were placed individually into glass cylinders (height: 23 cm, diameter: 20 cm) filled up to two-thirds with water at ∼24°C for 6 min. Automated scoring was done using the automated X’PERT FST software (Bioseb, Vitrolles, France).

### 2.4. Intracerebral in vivo microdialysis

Each mouse was anesthetized with chloral hydrate (400 mg/kg, i.p.) and implanted with microdialysis probes (CMA7 model, Carnegie Medicine, Stockholm, Sweden), two probes in the medial prefrontal cortex (right and left sides of the mPFC) and one in the dorsal raphe nucleus (DRN). Stereotaxic coordinates were as follows in mm from bregma: mPFC : A= + 2.2, L= ± 0.2, V= - 3.4; DRN (with an angle of 15°)/ A= - 4.5, L= ± 1.2, V= - 4.7; (A, anterior; L, lateral; and V, ventral) (Nguyen et al., 2013; Pham et al., 2018a and 2018b). On the same day, after awakening, mice received an acute dose of NBQX or muscimol 30 min before mPFC ketamine injection. On the next day, ∼24h after ketamine injection, the probes were continuously perfused with an artificial cerebrospinal fluid (aCSF, composition in mmol/L: NaCl 147, KCl 3.5, CaCl_2_ 1.26, NaH_2_PO_4_ 1.0, pH 7.4 ± 0.2) at a flow rate of 1.0 µl/min and 0.5 µl/min in the mPFC and DRN, respectively, using CMA/100 pump (Carnegie Medicine, Stockholm, Sweden), while mice were awake and freely moving in their cage. One hour after the start of aCSF perfusion stabilization period, four fractions were collected (one every 25 min) to measure the basal extracellular levels of 5-HT, glutamate and GABA (5-HT_ext_, Glu_ext_, GABA_ext_) in the mPFC as previously described (Pham et al., 2018a; Defaix et al., 2018). The lower limits of quantification (LLOQ) were ∼0.5 fmol/sample, ∼1.25 ng/ml and ∼0.6 ng/ml for 5-HT, glutamate and GABA, respectively. AUC values (% of baseline) were also calculated as previously described (Nguyen et al., 2013). At the end of the experiments, localization of microdialysis probes was verified histologically (Bert et al., 2004).

In the DHK protocol, after measurements of basal neurotransmitter levels, a 5 mM dose of DHK was perfused for 150 min in the mPFC (based on the study of Gasull-Camos et al., 2017), then five samples were collected. The FST was performed at the 3^rd^ sample (Figure 3A). After the perfusion, three other samples were collected to re-establish the baseline level of 5-HT_ext_, Glu_ext_, and GABA_ext_.

### 2.5. Statistics

All experimental results are given as the mean ± SEM. Data were analyzed using Prism 6 software (GraphPad, San Diego, CA, USA). A two-way ANOVA with pre-treatment (Vehicle vs NBQX or muscimol or GLT-1 inhibitor) and treatment (Vehicle vs ketamine) factors was used followed by Bonferroni post hoc test. A one-way ANOVA was also used to compare Vehicle vs DHK-treated mice, followed by Fisher’s PLSD post hoc test. Statistical significance was set at p ≤ 0.05.

## 3. Results

### 3.1. AMPA-R antagonist NBQX-pre-treatment into the DRN blocks the sustained antidepressant-like activity of ketamine and its releasing effects on 5-HT-glutamate-GABA in the mPFC

The microinfusion protocol of AMPA-R antagonist NBQX (0.25 nmol) in the DRN 30 minutes before intra-mPFC injection of ketamine (1 nmol/side) is described in Fig. 1A. A 2-way ANOVA revealed a significant main effect of ketamine [F(1,15) = 3.98, p = 0.05], no effects of NBQX in the FST [F(1,15) = 1.12, p = 0.30], and a significant interaction between these factors [F(1,15) = 10.1, p<0.01]. Post-hoc analyses indicated that ketamine significantly increased the swimming duration in the FST as compared to vehicle-treated mice (**p<0.01) (Fig. 1B). The swimming duration in the NBQX-ketamine treated group did not differ from the vehicle-treated group (p = 0.53). Thus, NBQX blocked the antidepressant-like activity of ketamine in the FST.

The analyses of 5-HT_ext_ in the mPFC indicated a significant main effect of ketamine [F(1,29) = 5.70, p<0.05], but did not show effects of NBQX [F(1,29) = 2.40, p = 0.13] and no interaction between these two factors [F(1,29) = 0.32, p = 0.57]. Post-hoc analyses indicated that ketamine significantly increased AUC values of 5-HT_ext_ as compared to vehicle-treated mice (*p<0.05) (Fig. 1F). 5-HT release in the NBQX-ketamine group did not differ from the vehicle-treated mice (p = 0.55). Thus, NBQX blunted ketamine-induced release of 5-HT in the mPFC.

The analyses of Glu_ext_ in the mPFC indicated a significant main effect of NBQX [F(1,28) = 5.85, p<0.05], and ketamine [F(1,28) = 6.08, p<0.05], but no interaction between these factors [F(1,28) = 0.17, p = 0.68]. Post-hoc analyses indicated that ketamine significantly increased AUC values of Glu_ext_ as compared to vehicle-treated mice (*p<0.05) (Fig. 1F). Glutamate release in the NBQX-ketamine group did not differ from the vehicle-treated mice (p = 0.97). Thus, NBQX blunted ketamine-induced release of glutamate in the mPFC (#p = 0.05).

The analyses of GABA_ext_ in the mPFC indicated a significant main effect of NBQX [F(1,28) = 4.07, p = 0.05], and ketamine [F(1,28) = 4.36, p<0.05], but no interaction between these two factors [F(1,28) = 1.23, p = 0.27]. Post-hoc analyses indicated that the ketamine increased GABA release in the mPFC as compared to vehicle-treated mice (*p<0.05) (Fig. 1F). GABA release in the NBQX-ketamine group did not differ from the vehicle-treated mice (p = 0.97). NBQX itself significantly increased AUC values of GABA_ext_ as compared to vehicle-treated mice (*p<0.05) (Fig. 2F).

Overall, at t24hr after treatment, ketamine alone increased swimming duration in the FST (Fig. 1B) and increased AUC values of 5-HT_ext_, Glu_ext_ and GABA_ext_ in the mPFC by 159%, 168% and 162%, respectively (Fig. 1C to 1F) compared to the vehicle-treated group. Intra-DRN NBQX prevented the effects of intra-mPFC ketamine injection on the swimming duration in the FST (Fig. 1B), and blunted the effects of ketamine on mPFC 5-HT_ext_, Glu_ext_ (Fig. 1F). By contrast, intra-DRN NBQX had no effects on ketamine-induced increase in cortical GABA_ext_ since ketamine-induced GABA release in the mPFC persisted following blockade of DRN AMPA-R (Fig. 1F). Thus, activation of DRN AMPA-R exerts a key control on ketamine-induced cortical 5-HT/glutamate release linked to its antidepressant-like activity in mice.

### 3.2. GABA_A_-R agonist muscimol pre-treatment in the mPFC decreases antidepressant-like activity and 5-HT releasing property of ketamine

To determine the influence of neuronal silencing on ketamine responses, muscimol was infused into the mPFC 30 minutes before intra-mPFC ketamine injection (see Fig. 2A, design of the protocol). Neurochemical and behavioral responses were assessed at t24hr post-injections to avoid the acute effects of drug treatments. A 2-way ANOVA revealed a significant main effect of muscimol in the FST [F(1,15) = 4.039, p = 0.05], ketamine [F(1,15) = 9.97, p<0.01], and interaction between these factors [F(1,15) = 5.72, p<0.05]. Post-hoc analyses indicated that ketamine significantly increased the swimming duration in the FST as compared to vehicle-treated mice (*p<0.05) as well as to muscimol-ketamine treated mice (#p<0.05) (Fig. 2B). The swimming duration in this later group did not differ from the vehicle-treated group (p>0.99). Thus, muscimol blocked the antidepressant-like activity of ketamine in the FST.

The analyses of 5-HT_ext_ in the mPFC indicated a significant main effect of ketamine [F(1,30) = 6.36, p<0.01], and interaction between muscimol and ketamine [F(1,30) = 6.97, p = 0.01], but did not show effects of muscimol [F(1,30) = 2.14, p = 0.15]. Post-hoc analyses indicated that ketamine significantly increased AUC values of 5-HT_ext_ as compared to vehicle-treated mice (**p<0.01) as well as to muscimol-pretreated mice (##p<0.01) (Fig. 2F). Thus, muscimol blocked ketamine-induced release of 5-HT in the mPFC.

The analyses of Glu_ext_ in the mPFC indicated a significant main effect of muscimol [F(1,30) = 5.57, p<0.05], but did not show effects of ketamine [F(1,30) = 0.28, p = 0.60] and no interaction between these two factors [F(1,30) = 2.47, p = 0.12]. Post-hoc analyses indicated that muscimol itself significantly increased AUC values of Glu_ext_ as compared to vehicle-treated mice (*p<0.05) (Fig. 2F). Thus, similar glutamate release occurred in ketamine-treated groups whether muscimol was perfused or not in the mPFC (p = 0.58).

The analyses of GABA_ext_ in the mPFC indicated a significant main effect of muscimol [F(1,30) = 4.69, p<0.05], and ketamine [F(1,30) = 3.44, p<0.05], but no interaction between these factors [F(1,30) = 0.187, p = 0.66]. Post-hoc analyses indicated that the combined perfusion of muscimol and ketamine increased GABA release in the mPFC as compared to vehicle-treated mice (p<0.05) (Fig. 2F).

Interestingly, a trend toward increased mPFC Glu_ext_ and GABA_ext_ was observed (AUC values increased by 144% (p=0.467), and 150% (p=0.11), respectively compared to the vehicle-treated group) (Fig. 2D to 2F). It constrasts with **ketamine-induced elevations in GABA and Glutamate that were reported in Fig. 1 or previously (Pham et al., 2018).** The complexity of the protocol combining the FST with microdialysis at t24h may explain this difference.

Overall, activation of mPFC GABA_A_-R by muscimol mainly controls ketamine responses on two serotonergic parameters, cortical 5-HT_ext_ and swimming duration. It suggests that the release of endogenous GABA by inhibitory interneurons located in the mPFC, and the subsequent activation of GABA_A_-R may limit ketamine responses. It also indicates that serotonin release in the mPFC is a key component of this response, and NMDA-R blockade is not sufficient to produce an antidepressant response (Fuchikami et al., 2015).

### 3.3 Intra-mPFC DHK, a glial glutamate transporter GLT-1 inhibitor, mimics the sustained antidepressant-like effect and neurochemical effects of ketamine (mPFC 5-HT/Glutamate/GABA release)

The glial GLT-1 glutamate transporter is mainly responsible for cortical glutamate reuptake (see Fig. 3A, design of the protocol).

Unpaired *t-*tests showed a statistically significance after of intra-mPFC DHK injection among the two groups (Fig. 3) in the FST (t = 2.68, *p<0.05, Fig. 3B), 5-HT_ext_ (t = 4.97, **p<0.01, Fig. 3C, 3F), Glu_ext_ (t = 2.83, *p<0.05, Fig. 3D, 3G) and GABA_ext_ (t = 4.23, **p<0.01, Fig. 3E, 3H) in the mPFC. Thus, the selective GLT-1 inhibitor induced a sustained antidepressant-like activity at t24h in the FST. This response appears to be mediated by increases in the three neurotransmitters studied. Overall, these data are similar to those described at t24h following a single intra-mPFC administration of ketamine.

## 4. Discussion

Here, a pharmacological approach was carried out to study the specific role of the mPFC-DRN circuit and excitatory/inhibitory balance in ketamine-induced antidepressant-like activity in BALBc/J mice. Our study assessed consequences of blocking DRN AMPA-R or mPFC GABA_A_-R on ketamine behavioral response associated with mPFC neurotransmitter release. Ketamine was injected locally into the mPFC, a brain region known to be involved in its fast antidepressant activity. Such a local injection strategy facilitates the analysis of the role of AMPA-R and GABA_A_-R located in the circuit mPFC-DRN in behavioral and neurochemical responses of ketamine. We report that ketamine-induced increase in swimming duration in the FST at t24h was blocked by a pre-treatment with either NBQX (AMPA-R antagonist) intra-DRN or muscimol (GABA_A_-R agonist) intra-mPFC. These data underline the involvement of DRN AMPA-R and mPFC GABA_A_-R in modulating the sustained ketamine antidepressant-like activity in BALB/cJ mice. It suggests that the activation of DRN AMPA-R is necessary to facilitate the sustained ketamine antidepressant-like activity, while activation of mPFC GABA_A_-R may limit this response (see below).

AMPA-Rs are located on DRN 5-HT neurons and endogenous glutamate can activate this ionotropic glutamatergic receptor subtype (Gartside et al., 2007). The present results regarding intra-DRN NBQX injection agree with previous studies who described the blockade of the antidepressant-like effects of ketamine following a systemic NBQX administration 30 min prior to testing in rats (Koike et al., 2011; Koike & Chaki, 2014) and in mice (Fukumoto et al., 2016; Koike et al., 2011; Pham et al., 2018b). Thus, activation of AMPA-R in the DRN would participate in the sustained antidepressant-like activity of ketamine (at t24h) by increasing mPFC serotonergic and glutamatergic neurotransmission. Interestingly, NBQX blocked ketamine-induced increase in 5-HT_1B_ receptor binding and decrease in SERT binding in primates (Yamanaka et al., 2014). By contrast, DRN AMPA-R activation does not seem to influence ketamine-induced cortical GABA release in mice.

A loop of regulation has been described from the cortex to the DRN and *vice versa*. Infusion of S-AMPA into the infralimbic cortex produced a rapid antidepressant-like response associated with increases in glutamate and 5-HT in this brain region (Gasull-Camos et al., 2017). Acting on the other part of the loop using a direct AMPA-R activation in the DRN increased 5-HT and glutamate release in the mPFC (Nishitani et al., 2014). Moreover, the activation of 5-HT neurons in the DRN is regulated by the stimulation of AMPA receptors in the mPFC (Fukumoto et a., 2016). Thus, a prominent role of the cortex-DRN circuitry and activation of the ascending 5-HT pathways in mediating a sustained antidepressant response was pointed out (Gasull-Camos et al., 2018). Ketamine also transiently increases spontaneous AMPA receptor-mediated neurotransmission in the DRN (Llamosas et al., 2019). In addition, a subcutaneous administration of ketamine increased the prefrontal 5-HT levels in a dose-dependent manner, which was attenuated by local injection of AMPA-R antagonists into the DRN.

Note that NBQX alone increased mPFC GABA_ext_. Several factors could explain this effect. First, it could be due to the basal anxiety phenotype already described in BALB/cJ mice (Dulawa et al., 2004; Holick et al., 2008). Second, it is possible that the dose of NBQX (0.25 nmol/side) infused into the DRN was too high. Indeed, it was already shown that the decrease in the immobility duration in the FST induced by systemic administration of ketamine (30 mg/kg, i.p.) was blocked by a lower dose of NBQX (0.03 nmol/side into the mPFC) in C57BL/6J mice, while NBQX per se had no effects (Fukumoto et al., 2016). Similarly, NBQX (30 nmol into the DRN 10 min before 25 mg/kg, s.c. ketamine) attenuated ketamine-induced 5-HT release in rat mPFC, while NBQX per se increased mPFC 5-HT_ext_ (Nishitani et al., 2014). By contrast, when given alone in rats, higher dose of NBQX (300 µM into the mPFC) did not change 5-HT_ext_ and Glu_ext_ in the mPFC compared to controls (Lopez-Gil et al., 2007). In these different studies, however, NBQX and NMDA-R antagonists were administered 30 min before neurochemical and behavioral tests, not at t24hr as in the present study. Third, the effect of NBQX per se suggests that endogenous GABA levels are high under physiological conditions and DRN AMPA-Rs tonically control GABA release in the mPFC. A tonic activation of AMPA-R in the DRN by the endogenous glutamate may exert a negative feedback control on GABA release in the mPFC. It was shown that the antidepressant-like activity of ketamine requires the activation of raphe 5-HT neurons (Fukumoto et al., 2016). Reciprocally, the activity of mPFC glutamatergic pyramidal neurons is controlled by raphe 5-HT neurons (Puig et al., 2005; Gasull-Camos et al., 2014). Although this mPFC-DRN circuit seems to be important in ketamine responses (Hajos et al., 1998; Peyron et al., 1998; Vertes RP, 2004; Carreno et al., 2016), we cannot exclude a local effect of glutamate on 5-HT nerve terminals in the mPFC. Our experiment combining *in vivo* microdialysis and FST in the same mice provides, for the first time, more insights into how the blockade of DRN AMPA-R interacts with ketamine’s responses.

The main hypothesis regarding the indirect mechanism of action of ketamine is the disinhibition hypothesis explaining the increased burst of pyramidal glutamatergic neuron in the mPFC following the blockade of NMDA-R by ketamine (Miller et al., 2016). Parvalbumin-expressing interneurons provide inhibition of pyramidal glutamatergic neurons under physiological conditions. Ketamine selectively antagonizes these GABAergic interneurons to excitatory synapses leading to the loss of the tonic inhibition, thus increasing the burst of pyramidal glutamatergic neurons, leading to the release of mature Brain-Derived Neurotrophic Factor, which is required for the antidepressant-like activity of ketamine (Liu et al., 2012). However, this glutamate burst induces synaptic remodeling and resetting of glutamate and GABA systems (Duman et al., 2019). An increase in GABA-release as found here in the mPFC, appears to contradict this main hypothesis. It is proposed that the resulting burst of glutamate may lead to BDNF release and the sustained release of both glutamate and GABA (Duman et al., 2019). Such an upregulated GABA synaptic function, as presented here in mice, is in line with deficits of GABA measured in depressed patients (Sanacora et al., 2004).

A direct infusion of muscimol into the mPFC abolished the sustained antidepressant-like effects of ketamine in rats (Fuchikami et al., 2015), suggesting an interaction between NMDA and GABA_A_ receptors specifically in the mPFC. However, muscimol *per se* had no significant effect on the swimming duration when the time point analysis was t24hr (Fuchikami et al., 2015), but reduced immobility in the FST when the analysis was performed immediately after intra-mPFC muscimol administration (Slattery et al., 2011). In rat cortical neurons in culture, muscimol increased inhibitory neurotransmission by opening GABA-Cl(-)-channel, thus allowing inward Cl^-^ fluxes in cortical neurons and increase in basal glutamate release by potentiating intracellular Ca^2+^ influx. This effect is reversed by bicuculline suggesting a role of GABA_A_-R agonism in the control of neuronal excitation (Herrero et al., 1999]. Here, this excessive increase in mPFC Glu_ext_ by muscimol could cause an excitotoxicity and prevent beneficial stimulation of serotonergic neurons for antidepressant-like response of ketamine.

Surprisingly here muscimol *per se* enhanced mPFC Glu_ext_ (AUC values = 244% vs vehicle) in BALB/cJ mice indicating that GABA_A_-R was tonically activated by endogenous GABA levels under our experimental conditions, i.e., at t24h. A combination of a low dose of muscimol (0.1 mg/kg, i.p., 30 min prior to ketamine) with a very low dose of ketamine (0.1 mg/kg, i.p.) and tested at 30 minutes post-injection produced a synergistic antidepressant-like activity in the mouse tail suspension test (Rosa et al., 2016). Thus, the time point of observation is crucial because an acute effect of muscimol could result in increasing locomotion and decreasing immobility in these behavioral tests, while at 24h post-injection muscimol-induced inactivation of the mPFC by activating GABA_A_-R could result in a hindrance of ketamine’s effects.

The question then arises whether or not the present findings in mice could also occur following a co-administration of BZD and ketamine in patients. Few clinical studies suggest that co-administration of drugs interfering with GABAergic neurotransmission may hamper optimal ketamine response (Frye et al., 2015; Andrade, 2017). In a clinical trial carried out in TRD-BZD users, the benzodiazepine interfered with the antidepressant response produced by the first ketamine administration, but not subsequent treatments (Albott et al., 2017). Consistent with the suppression of excitatory glutamatergic networks by ketamine, a GABA_A_-R agonism (as occurs with a BZD) increased inhibitory tone of interneurons, thereby decreasing the therapeutic efficacy of ketamine.

In rodents, diazepam inhibited ketamine-induced hyperlocomotion (30 mg/kg) in mice (Irifune et al., 1998). This effect may be related to the BZD ability to suppress the activation of dopamine neurons in the nucleus accumbens and striatum. Moreover, the effects of intra-mPFC injections of GABAergic drugs on ketamine-induced amnesia have been studied in mice (Farahmandfar et al., 2017). Muscimol pre-treatment inhibited ketamine-induced memory formation (5 mg/kg i.p.), suggesting that GABAergic system is involved in ketamine-induced impairment of memory acquisition. Interestingly, treatment with agents activating GABA_A_-R such as diazepam reduced the severity of the psychotomimetic adverse effects of ketamine in rodents, and is used in human anesthesia for this purpose (Olney et al., 1991; Irifune et al., 2000). Diazepam was especially helpful in reducing the emergence delirium of ketamine (Fontenot et al., 1982).

The local administration of DHK into the mPFC mimics the sustained effects of ketamine by inhibiting the glutamate transporter GLT-1. DHK evoked an antidepressant-like activity at t24h associated with a robust increase in mPFC 5-HT_ext_, Glu_ext_ and GABA_ext_ in BALB/cJ mice. GLT-1 (or EAAT2), together with EAAT1, are two most abundant glutamatergic transporters in the forebrain (Gegelashvili et al., 2000). It is mainly located in glial cells and neurons and responsible for cortical glutamate reuptake (Gasull-Camos et al., 2017). In the chronic unpredictable stress model in rats, decreased levels of GLT-1 was observed in the hippocampus, and a single ketamine injection (10 mg/kg, i.p.) alleviated this abnormality (Liu et al., 2016). In our previous study, ketamine did not influence the function of glutamatergic transporters (see the zero-net-flux quantitative microdialysis experiment in Fig. 1G in Pham et al., 2018a), indicating that ketamine-induced increases in mPFC Glu_ext_ was of neuronal origin, rather than from astrocytes. Similar results have been obtained with a microinfusion of DHK in non-stressed rats, i.e., increases in swimming duration in the FST and increases in mPFC microdialysate Glu_ext_ and 5-HT_ext_ (Gasull-Camos et al., 2017). Thus, an acute increase in excitatory glutamate neurotransmission selectively in the mPFC triggers the sustained antidepressant-like activity of DHK in rodents. Using 5-HT synthesis inhibition, these authors also showed that these DHK responses are mediated by the activation of mPFC-raphe pathways, which then induced a fast increase in serotonergic activity (Gasull-Camos et al., 2017). The increase in Glu_ext_ in cortical synapses induced by GLT-1 blockade is likely the source of this neuronal excitation. However, by measuring GABA_ext_ here, we show that this phenomenon occurred with a constant ratio of Glu_ext_/GABA_ext_ (see a summary the behavioral and neurochemical effects of ketamine in Table 1) indicating that a homeostatic regulation of the excitation/inhibition balance was maintained in the mPFC, as hypothesized by Bernhard Luscher’s group (Ren et al., 2016). Similar effects of DHK and ketamine suggest a rapid neural-glial adaptation involved in the sustained antidepressant effects of ketamine. Whether such a neuronal/glial relationship supports the rapid (1-2 hrs post dose) and/or the sustained (at t24h) antidepressant-like activity of ketamine requests further investigations.

**Table 1.**
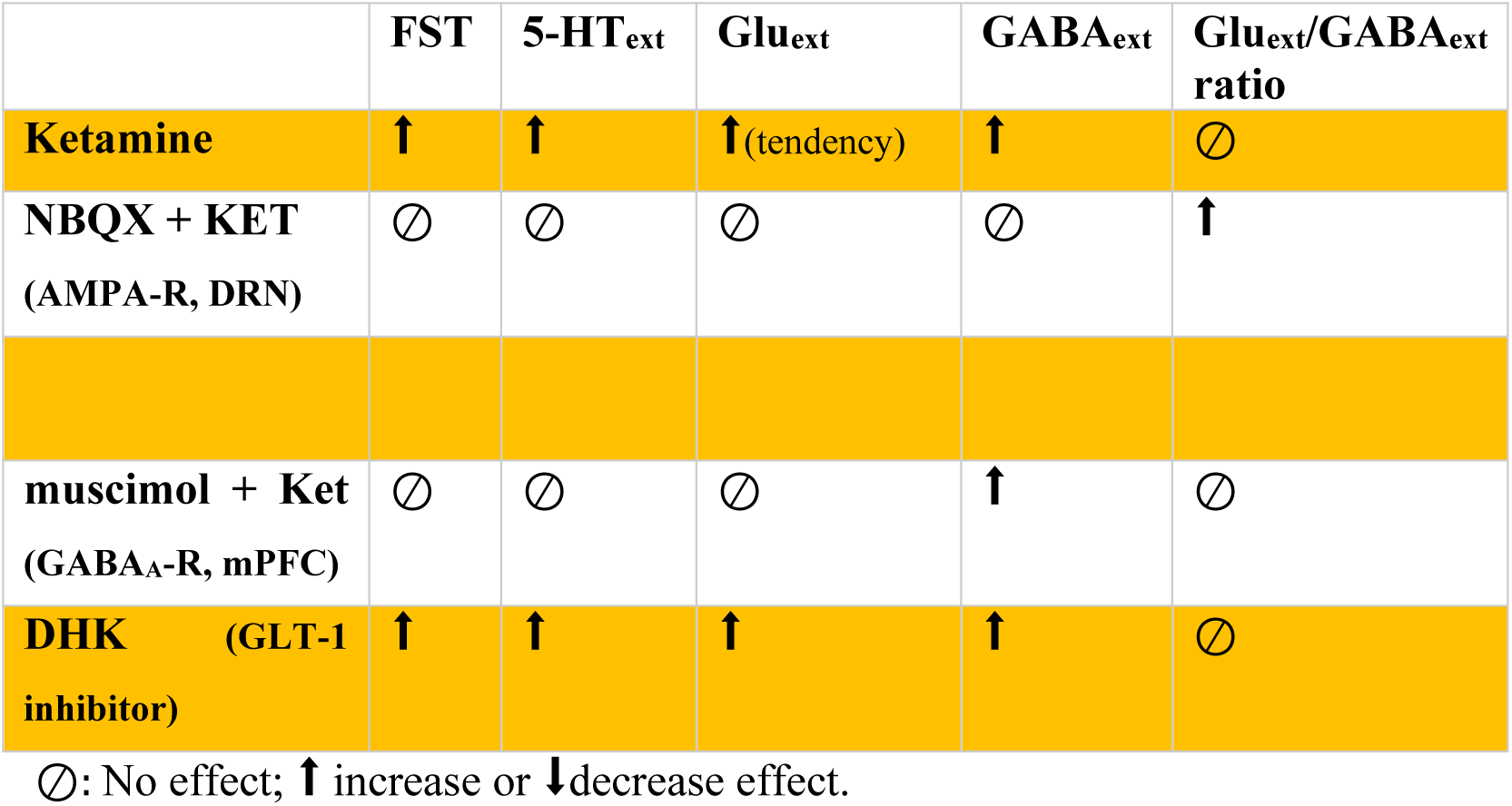
Summary of behavioral and neurochemical effects (in the mPFC) obtained in the present study

|  | **FST** | **5-HT<sub>ext</sub>** | **Glu<sub>ext</sub>** | **GABA<sub>ext</sub>** | **Glu<sub>ext</sub>/GABA<sub>ext</sub>**<br>**ratio** |
| --- | --- | --- | --- | --- | --- |
| **Ketamine** | ↑ | ↑ | ↑<sub>(tendency)</sub> | ↑ | ∅ |
| **NBQX + KET**<br>(AMPA-R, DRN) | ∅ | ∅ | ∅ | ∅ | ↑ |
| **muscimol + Ket**<br>(GABA<sub>A</sub>-R, mPFC) | ∅ | ∅ | ∅ | ↑ | ∅ |
| **DHK** (GLT-1<br>inhibitor) | ↑ | ↑ | ↑ | ↑ | ∅ |

∅: No effect; ↑ increase or ↓ decrease effect.

In conclusion, we carried out here a pre-clinical approach to study the role of mPFC-DRN circuit and AMPA-R/DRN and GABA_A_-R/mPFC in the sustained antidepressant-like activity of ketamine. An acute increase in glutamate neurotransmission, either with ketamine and DHK (a GLT-1 glutamatergic transporter blocker) induced similar behavioral and neurochemical effects. This sustained antidepressant-like activity in mice is likely mediated by glutamate and 5-HT release in the mPFC in BALB/cJ mice (Cryan et al., 2002). Ketamine-induced GABA release in the mPFC persisted following blockade of DRN AMPA-R or activation of mPFC GABA_A_-R. The results of the present study could contribute to a further understanding of the cellular mechanisms underlying ketamine antidepressant-like activity.

Anxiety frequently coexists with depression and adding benzodiazepines to antidepressant treatment is common practice to treat people with major depression. The 2019 updated version of a Cochrane Review concluded that the combined antidepressant (TCAs, SSRIs) plus BZD therapy was more effective than antidepressants alone in improving depression severity, response in depression and remission in depression in the early phase (Ogawa et al., 2019). The present results should encourage clinicians to test for the concomitant prescription of ketamine and BZD to see whether its sustained antidepressant activity is maintained in TRD patients.

## Abbreviations

5-HT: Serotonin
aCSF: Artificial cerebrospinal fluid
AD: Antidepressant drug
AMPA: α-amino-3-hydroxy-5-methylisoxazole-4-propionic acid
AMPA-R: AMPA receptor
AUC: Area under the curve
BZD: Benzodiazepine
DHK: Dihydrokainic acid
DRN: Dosal raphe nuclei
EAAT2: Excitatory amino acid transporter 2
FST: Forced swim test
GABA: γ-Aminobutyric acid
GABA_A_: γ-Aminobutyric acid type A
GABA_A_-R: GABA_A_ receptor
GLT-1: Glutamate transporter 1
GLUT-1: Glutamate transporter
KET: Ketamine
LLOQ: Lower limits of quantification
mPFC: Medial prefrontal cortex
NMDA N-: methyl-D-aspartate
NMDA-R: NMDA receptor
SEM: Standard error of the mean
SERT: Serotonin transporter
SSRI: Selective serotonin reuptake inhibitor
TRD: Treatment-resistant depressed

## Acknowledgements

Dr. Thu Ha Pham was supported by a fellowship from the “*Ecole Doctorale 569: Innovation Therapeutique*”. The authors would like to thank colleagues from the animal care facility of SFR-UMRS “Institut Paris Saclay d’Innovation Thérapeutique” of University Paris-Sud for their technical assistance. A special thanks to Audrey Solgadi and Pierre Chaminade from the platform “*Service d’Analyse des Médicaments et Métabolites (*SAMM) » of the Faculty of Pharmacie to set up the mass spectrometry equipment. Our team UMR-S 1178, Inserm, Univ Paris-Sud/Paris-Saclay provided the necessary resources to perform this study.

## Authors’ contribution

All authors contributed to the conception and design of the study; THP, TMLN, CD, LT and DJD contributed to the acquisition of data. THP, TMLN and AMG wrote the manuscript. All the authors contributed to analysis of data, drafting the article for key intellectual content.

